# Working memory demands modulate memory brain state engagement

**DOI:** 10.64898/2026.04.14.718495

**Authors:** DT Nguyen, Nicole M. Long

## Abstract

The extent to which attention and memory processes rely on shared as opposed to distinct mechanisms is critical for understanding the role of both processes in cognition. As working memory sits at the intersection of external, perceptual input and long-term internal storage, it provides the ideal testbed for investigating overlaps between attention and memory. We hypothesize that memory brain states, whole-brain activity patterns that support long-term memory encoding and retrieval, map onto the external/internal axis of attention. Specifically, we hypothesize that external attention, focusing on sensory information, recruits the encoding state and internal attention, focusing on stored information, recruits the retrieval state. To test this hypothesis, we conducted a scalp electroencephalography study in which participants engaged in a working memory paradigm with and without maintenance demands. We used an independently validated cross-study multivariate pattern classifier to measure memory brain state engagement during change and target detection tasks. We find that the encoding state is recruited for stimulus presentation during both tasks, whereas the retrieval state is selectively recruited during the delay of the change detection task. Together, these results suggest that memory states map onto the external/internal axis of attention to support working memory, long-term memory, and cognition more broadly.

## Introduction

Upon arrival at the airport, you try to form a memory of the current moment so that you can find your car on your return. You direct your resources to sensory information in the world around you – an alphanumeric sign on a concrete column, a yellow painted number under your car. When you return from your trip, you direct your resources to the stored information to recover those no longer present sensory details. Thus memory – in particular, the processes of encoding and retrieving experiences – is tightly linked to attention, or how limited capacity resources are directed (Nobre, 2018). Memory encoding and memory retrieval constitute neurally distinct brain states that trade off – meaning that encoding and retrieval cannot occur simultaneously – and this tradeoff impacts behavior (Long & Kuhl, 2019). Memory brain states may reflect processes specific to episodic, long-term memory, that is, the formation and retrieval of events situated within a particular spatiotemporal context (Tulving, 1983). However, growing research suggests that memory brain states may extend beyond episodic memory and instead map onto the external/internal axis of attention (Chun, Golomb, & Turk-Browne, 2011). External attention constitutes focusing on sensory, environmental information, whereas internal attention constitutes focusing on mental processes and representations. Establishing a link between memory brain states and attention is critical for determining how both memory and attention impact cognition. As working memory (WM) sits at the intersection of external and internal attention, it is well suited for assessing such a link. The aim of this study is to directly connect memory brain states with the external/internal axis of attention via a WM paradigm with and without memory demands.

Establishing the degree to which memory brain states reflect how attention is oriented is critical both to the specific goal to identify the mechanisms that support memory and attention and to the broad goal of understanding how brain states impact cognition. Although often considered in isolation, memory and attention have a strong degree of overlap, both conceptually and neurally (Chun & Turk-Browne, 2007; Cabeza, Ciaramelli, Olson, & Moscovitch, 2008; Sestieri, Shulman, & Corbetta, 2010; Long, Kuhl, & Chun, 2018; Fischer, Moscovitch, & Alain, 2020). A link between memory brain states and external/internal attention would constrain models of cognition (Logan, Cox, Annis, & Lindsey, 2021) and enable online, real-time identification of when and how specific memory and attentional processes are engaged. Furthermore, brain states, or whole brain activity/connectivity patterns (Harris & Thiele, 2011), modulate how stimuli are processed (Long & Kuhl, 2021), impact behavior (Harris & Thiele, 2011; Kay & Frank, 2019), and influence the efficacy of brain stimulation efforts (Bradley, Nydam, Dux, & Mattingley, 2022). Characterizing the factors that induce memory brain state engagement will further understanding of how orienting attention externally vs. internally modulates behavior.

Growing evidence suggests that the retrieval state tracks internal attention (Logan et al., 2021; Long, 2023; Cleary et al., 2025; Han & Long, 2025; Bair & Long, 2026). The retrieval state is recruited under explicit demands to retrieve episodic memories (Smith & Long, 2024) as originally proposed (Tulving, 1983). However, recent work has shown that the retrieval state is also engaged when participants retrieve from semantic memory (Bair & Long, 2026) and selectively attend to particular spatial locations (Long, 2023). The retrieval state is characterized by a sustained pattern of voltage topography that is thought to reflect engagement of the default mode network (DMN; Raichle et al., 2001; Britz, Van De Ville, & Michel, 2010; Custo et al., 2017; Hong, Moore, Smith, & Long, 2023). The DMN supports episodic and semantic memory retrieval (Binder & Desai, 2011; H. Kim, 2016) and more broadly, internal mentation (Buckner & DiNicola, 2019). In tandem, scalp electroencephalography (EEG) studies have revealed an overlap in the spectral signals that support selection from both perception and memory (Woodman, Wang, Sutterer, Reinhart, & Fukuda, 2022). Specifically, spectral power in the alpha band (10-14 Hz) is modulated both by focusing attention internally (Ray & Cole, 1985; Cooper, Croft, Dominey, Burgess, & Gruzelier, 2003; Palva & Palva, 2007; Villena-González, López, & Rodríguez, 2016), as well as by long-term memory retrieval (e.g. Martín-Buro, Wimber, Henson, & Staresina, 2020). Finally, behavioral performance in an episodic version of the flanker task (Eriksen & Eriksen, 1974) suggests that retrieval from short and long-term memory requires inward focused attention (Logan, Lilburn, Ulrich, Weeks, & Koo, 2026). Taken together, these findings suggest that the retrieval state is not specific to long-term episodic remembering, but tracks the degree to which the mind’s eye is turned inward.

As WM sits at the intersection between external and internal attention (Chun et al., 2011), it serves as the ideal testbed for linking memory states to attention. WM enables the maintenance and manipulation of information in the absence of sensory input (Baddeley & Hitch, 1974). Like attention more broadly (Marois & Ivanoff, 2005), WM has capacity limits whereby only a small amount of information can be maintained (Luck & Vogel, 1997; Cowan, 2010; Luck & Vogel, 2013; Oberauer, 2019). Alpha power tracks both perceptual and WM load (e.g. Kornblith, Buschman, & Miller, 2016), though there is some variability as to whether maintenance-related alpha power increases (Jensen, Gelfand, Kounios, & Lisman, 2002; Heinrichs-Graham & Wilson, 2015; Schroeder, Ball, & Busch, 2018; Hu et al., 2019) or decreases (Bae & Luck, 2018; Erickson, Smith, Albrecht, & Silverstein, 2019; Chen, van Ede, & Kuo, 2022) as a function of load. WM paradigms tap both external and internal attention, with external attention recruited during the encoding or display interval and internal attention recruited during the delay interval (Vo et al., 2021). Display interval alpha power decreases (Cona et al., 2020) along with delay interval alpha power increases (Jensen et al., 2002) are consistent with the broader finding that alpha dissociates external vs. internal attention. To the extent that memory brain states selectively reflect long-term episodic remembering, there should be no memory state modulation within the context of a WM paradigm. Alternatively, to the extent that memory brain states track attentional orienting, the shifting demands within a WM task should induce memory brain state fluctuations.

Our central hypothesis is that memory states map onto the external/internal axis of attention, whereby external attention recruits the encoding state and internal attention recruits the retrieval state. We have two specific hypotheses. First, we hypothesize that the encoding state will be recruited during initial stimulus presentation regardless of memory demands. Alternatively, the encoding state will be selectively recruited during stimulus presentation in response to top-down demands to encode the stimulus display. Second, we hypothesize that WM maintenance will specifically recruit the retrieval state. Alternatively, the retrieval state will be selectively recruited in response to top-down demands to retrieve a past experience. To test our hypothesis, we conducted a scalp EEG study in which participants performed either a target detection task or a change detection task. We used cross-study multivariate pattern analyses (MVPA) to assess how encoding and retrieval states are engaged by fluctuating WM demands. To the extent that memory brain states reflect external and international attention, these states should be differentially recruited across the phases of this task.

## Materials and Methods

### Participants

Forty participants (33 female; age range = 18-35 years, mean age = 19.88 years) from the University of Virginia community participated. Our sample size was determined *a priori* based on our prior work and is described in the pre-registration report of this study (https://osf.io/rcwk3). All participants had normal or corrected-to-normal vision. Informed consent was obtained in accordance with the University of Virginia Institutional Review Board for Social and Behavioral Research, and participants were compensated for their time. One participant was excluded from the final dataset due to poor task performance (only responded to 3% of trials). Thus, data are reported for the remaining 39 participants.

### Experimental Design and Statistical Analysis

#### Procedure and Design

##### General overview

Stimuli were colored squares presented on a grey background with eight possible colors (Figure 1). Participants viewed 1, 2, 4, or 6 squares per trial. Square locations were random across the screen, constrained to one of twenty-four locations on a 4 by 6 grid (not shown to participants) with a small jitter. To manipulate memory demands, trials were divided into perception blocks and memory blocks. Each of the twelve runs contained two blocks, memory and perception, and block order was random within run. Each block contained 24 trials, two response options, and participants were instructed to respond as quickly and accurately as possible. Each block began with a get ready screen indicating the current block and response mappings (3000 ms) followed by a 1000 ms inter-stimulus interval (ISI). Each trial began with a 500 ms display phase in which 1, 2, 4, or 6 squares were presented. The display phase was immediately followed by the delay phase in which a fixation cross was presented for 2000 ms. The delay phase was followed by the response phase. Participants pressed one of two response keys which was counterbalanced across participants. Responses were required within 1000 ms of response phase onset. The response phase was followed by a 1500 ms ISI.

**Figure 1.**
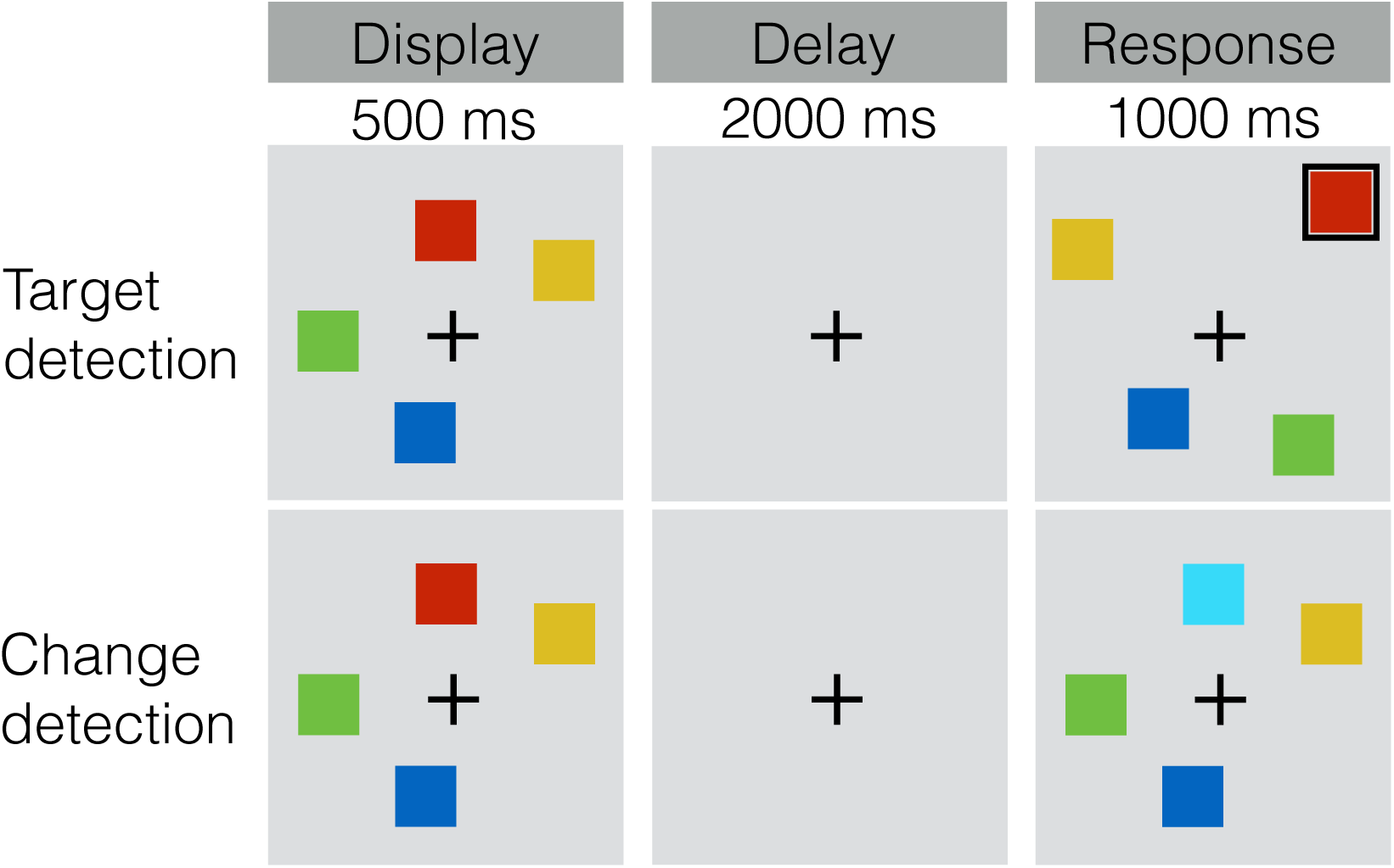
Task design. In both the target detection (perception) task and the change detection (memory) task, participants saw one, two, four, or six squares on the screen for 500 ms. Each display phase was followed by a 2000 ms delay phase. During the response phase, participants saw the same squares either in new locations with a white or black outline around one square (target detection) or the same squares in the same locations (change detection). On half of the change detection trials one of the squares changed to a complementary color as shown above. Participants’ task was to respond as to whether there was a white/black outline in the target detection task or to respond as to whether one of the squares had changed color in the change detection task. Participants had 1000 ms to make their response.

##### Perception task

In the perception blocks, participants performed a target detection task. All squares from the display phase were re-presented in new locations with either a black or white outline around one of the squares. The locations changed to minimize (and potentially eliminate) participants’ tendency to maintain information about the squares across the delay. All square colors were identical to the display phase. Participants were instructed to respond with either the *d* or *k* key indicating the outline color.

##### Memory task

In the memory blocks, participants performed a change detection task. All, or all but one, squares from the display phase were re-presented in the same locations. On half of the memory trials, one square changed color between the display and response phase. The color changed to the complementary color of a square presented during the display phase and was not utilized for any of the other squares in either the display or response phase (e.g. a red square would change to cyan and no other squares in the display would be cyan). Participants were instructed to respond with either the *d* or *k* key indicating whether any of the squares did or did not change color from the display phase.

### EEG data acquisition and preprocessing

EEG recordings were collected using a BrainVision system and an ActiCap equipped with 64 Ag/AgCl active electrodes positioned according to the extended 10-20 system. All electrodes were digitized at a sampling rate of 1000 Hz and referenced to electrode FCz. Offline, electrodes were later converted to an average reference. Impedances of all electrodes were kept below 50kΩ. Electrodes that demonstrated high impedance or poor contact with the scalp were excluded from the average reference but included in all other analysis steps. Bad electrodes were determined by voltage thresholding.

Custom Python codes were used to process the EEG data. We applied a high pass filter at 0.1 Hz, followed by a notch filter at 60 Hz and harmonics of 60 Hz to each participant’s raw EEG data. We then performed three preprocessing steps (Nolan, Whelan, & Reilly, 2010) to identify electrodes with severe artifacts. First, we calculated the mean correlation between each electrode and all other electrodes as electrodes should be moderately correlated with other electrodes due to volume conduction. We z-scored these means across electrodes and rejected electrodes with z-scores less than -3. Second, we calculated the variance for each electrode as electrodes with very high or low variance across a session are likely dominated by noise or have poor contact with the scalp. We then z-scored variance across electrodes and rejected electrodes with a *|z| >*= 3. Finally, we expect many electrical signals to be autocorrelated, but signals generated by the brain versus noise are likely to have different forms of autocorrelation. Therefore, we calculated the Hurst exponent, a measure of long-range autocorrelation, for each electrode and rejected electrodes with a *|z| >*= 3. Electrodes marked as bad by this procedure were excluded from the average re-reference. We then calculated the average voltage across all remaining electrodes at each time sample and re-referenced the data by subtracting the average voltage from the filtered EEG data. Although bad electrodes were excluded from the average re-reference calculation, all electrodes were included for every participant for all subsequent analyses. We used wavelet-enhanced independent component analysis (Castellanos & Makarov, 2006) to remove artifacts from eye blinks and saccades.

### EEG data analysis

We applied the Morlet wavelet transform (wave number 6) to the entire EEG time series across electrodes, for each of 46 logarithmically spaced frequencies (2-100 Hz; Long & Kahana, 2015). After log-transforming the power, we downsampled the data by taking a moving average across 100 ms time intervals from 500 ms preceding to 4000 ms following trial onset, and sliding the window every 25 ms, resulting in 177 time intervals (45 non-overlapping). Power values were then z-transformed by subtracting the mean and dividing by the standard deviation power. Mean and standard deviation power were calculated across all trials and across time points for each frequency.

### Pattern classification analyses

Pattern classification analyses were performed using penalized (L2) logistic regression, implemented via the sklearn linear model module in Python. “Memory state evidence” was a continuous value reflecting the logit-transformed probability that the classifier assigned a particular mnemonic label (encode, retrieve) to each trial. Memory state evidence served as a trial-specific, continuous measure of memory state information, which we used to assess how encoding and retrieval states were engaged by fluctuating working memory demands.

We developed and validated a cross-participant mnemonic state classifier based on already-collected data from participants (N = 143) who completed a mnemonic state task (Hong et al., 2023; Han & Long, 2025). In this paired-objects task, participants view two lists of images of common objects. All List 1 objects are new. All List 2 objects are also new, but are categorically associated to a List 1 object. For example, if List 1 contains an image of a bench, List 2 would contain an image of a different bench. During List 1, participants are instructed to encode each object. During List 2, however, each trial contains an instruction to either encode the current object (e.g., the new bench) or to retrieve the corresponding object from List 1 (e.g., the old bench). We conducted three stages of classification using the same methods as in our prior work (Long, 2023; Smith & Long, 2024; Wheelock & Long, 2024; Han & Long, 2025). First, we performed within-participant leave-one-run-out classification (penalty = 1) on patterns of spectral power across 63 electrodes and 46 frequencies to identify the participants who show robust classification accuracy. We used data-derived chance classification accuracy values to identify a subset of participants (N = 57) for whom true classification accuracy was at or above the 90th percentile of permuted chance values. Second, using leave-one-participant-out cross-validated classification (penalty = 0.0001), we obtained classification accuracy of 59.29% which is significantly above chance (*t* _56_ = 7.667, *p <* 0.0001), indicating that the cross-participant mnemonic state classifier can distinguish memory encoding and memory retrieval states. Finally, we applied the cross-participant mnemonic state classifier to assess memory state evidence across all phases and tasks in the current study.

### Statistical Analyses

We used repeated measures ANOVAs (rmANOVAs) and linear and logistic regression along with paired t-tests to assess the effect of set size on behavior in the perception and memory tasks. We used rmANOVAs to assess the impact of task (memory or perceptual) and time on memory state evidence. We used paired *t* -tests to assess task differences in memory state evidence for the display and delay phases. We used linear and logistic regression to test the extent to which memory state evidence predicts subsequent memory. We used linear regression to relate memory state evidence to set size. We used false discovery rate (FDR; Benjamini & Hochberg, 1995) to correct for multiple comparisons.

## Results

### Set size modulates target and change detection behavior

Our first goal was to assess the impact of set size on target detection reaction time (RT) in the perception task and change detection accuracy in the memory task. We expected to find slower RTs in the perception task and lower accuracy in the memory task as set size increases. Following our pre-registration, we conducted two regressions to predict behavior as a function of set size. For the perception task, we conducted a linear regression with set size as the predictor and RT on accurate perceptual trials as the dependent variable (Figure 2A). We found a significant effect of set size on RT (*β* = 2.914, *SD* = 3.631, *t* _38_ = 4.948, *p <* 0.0001, *d* = 0.8026). For the memory task, we conducted a logistic regression with set size as the predictor and accuracy on memory trials as the dependent variable (Figure 2B). We found a significant effect of set size on memory accuracy (*β* = -0.4088, *SD* = 0.1938, *t* _38_ = -13.00, *p <* 0.0001, *d* = 2.109). These results show that as set size increases, target detection RTs increase and change detection accuracy decreases.

**Figure 2.**
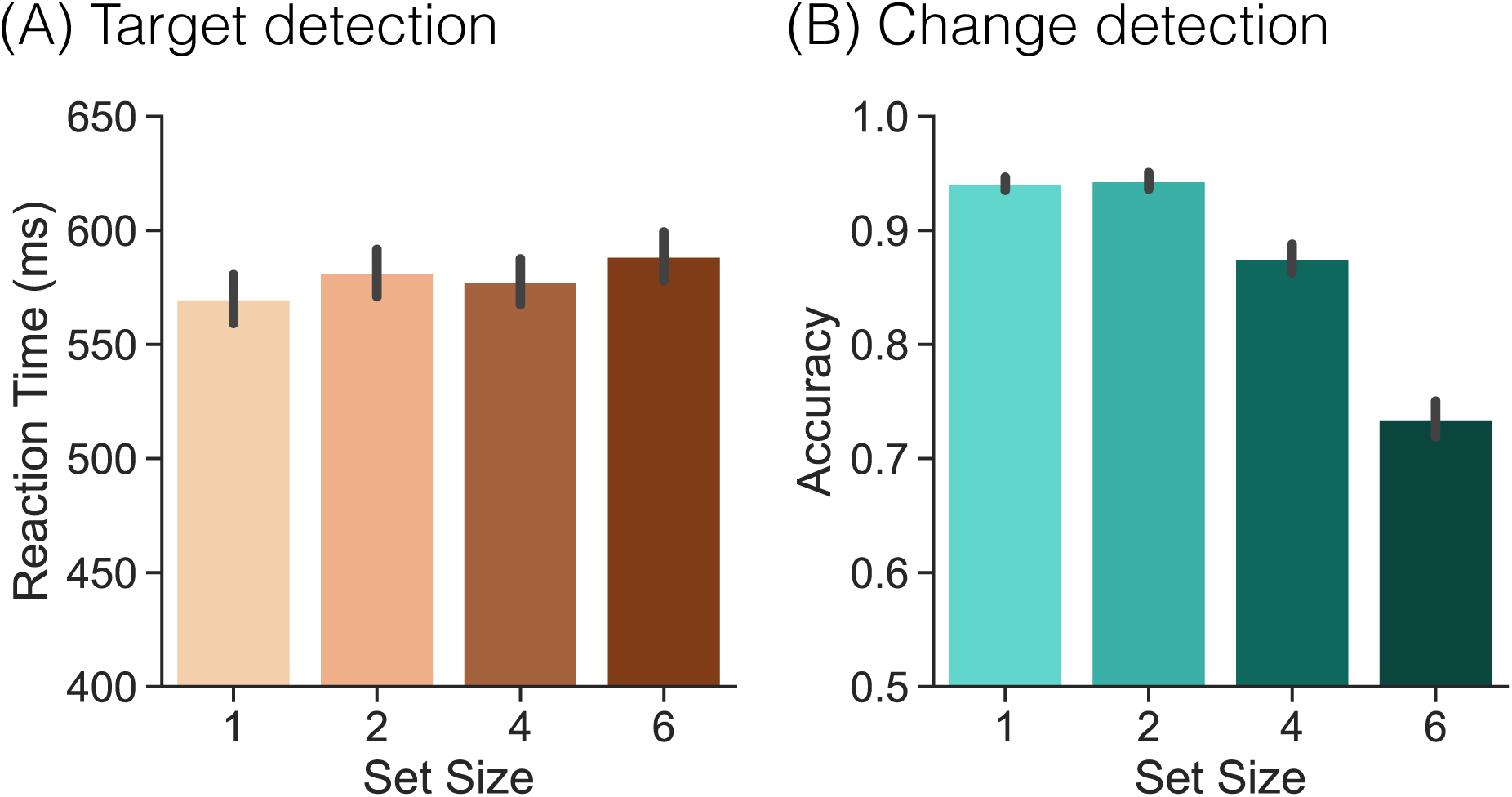
Influence of set size on behavior. (A) Target detection reaction times as a function of set size (1, 2, 4, 6). There is a significant increase in reaction time as a function of set size (*p <* 0.0001). (B) Change detection accuracy as a function of set size (1, 2, 4, 6). There is a significant decrease in change detection accuracy as a function of set size (*p <* 0.0001). Error bars denote standard error of the mean.

### Memory brain states are modulated by working memory demands

To generally assess the impact of dynamic WM demands – including the presumed external/internal attention demands – on memory brain states, we first measured memory state evidence over time across the entire trial separately for the perception and memory tasks (Figure 3). Following our pre-registration, we conducted a 2 *×* 35 repeated measures ANOVA (rmANOVA) with factors of task and time interval (100 ms intervals from 0 ms to 3500 ms following display onset). Stimulus-locked memory state evidence served as the dependent variable. We found a significant main effect of time (*F* _34,1292_ = 35.96, *p <* 0.0001, 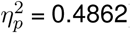) and task (*F* _1,38_ = 29.99, *p <* 0.0001, 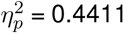). We found a significant interaction between task and time (*F* _34,1292_ = 18.10, *p <* 0.0001, 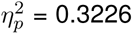). Thus, memory state engagement varies as a function of task and over time.

**Figure 3.**
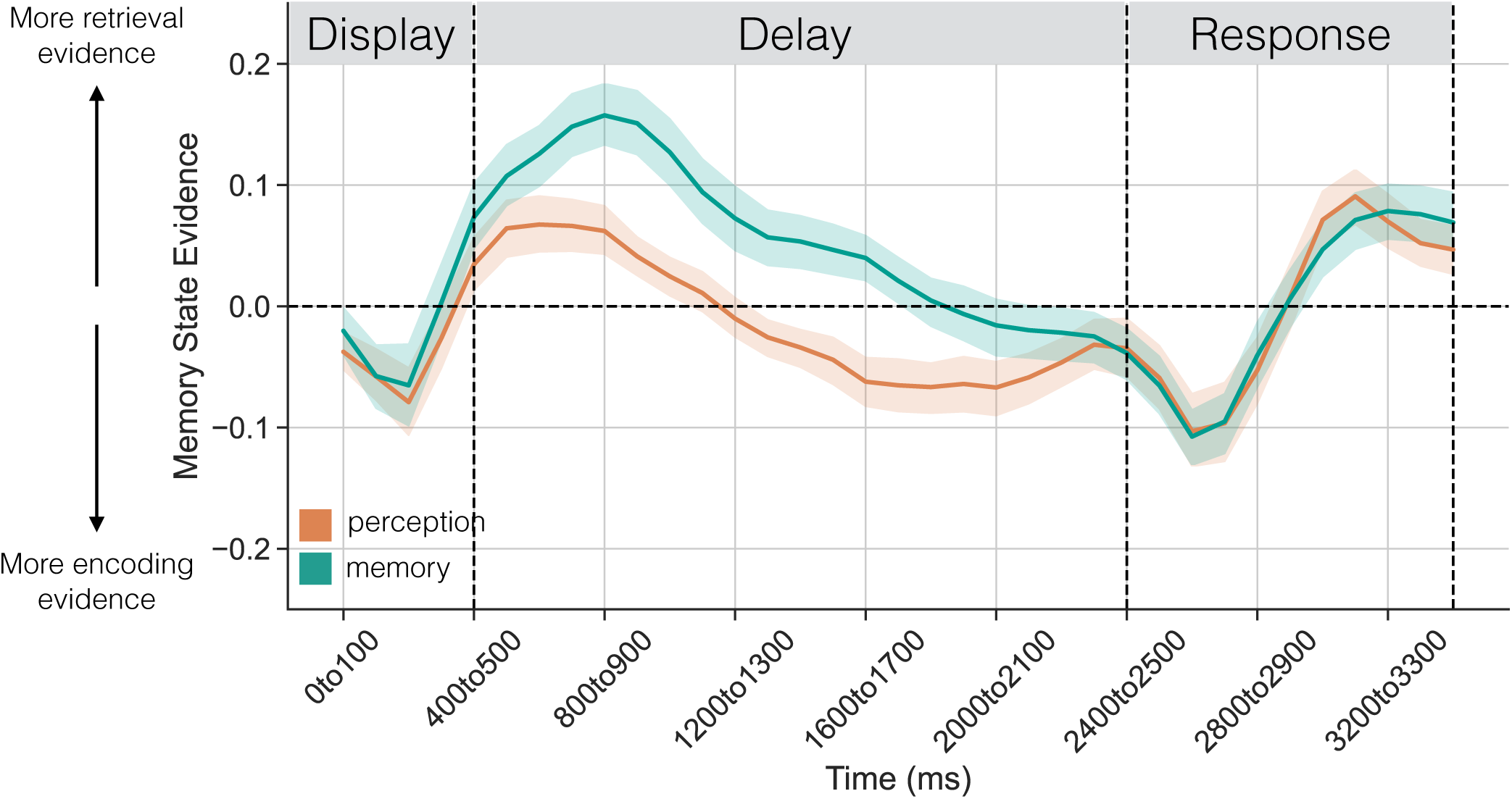
Memory state evidence over time. We assessed memory state evidence as a function of task (perception, orange; memory, teal) across display (0 to 500 ms), delay (500 to 2500 ms), and response (2500 to 3500 ms) phases. Positive values indicate greater evidence for retrieval; negative values indicate greater evidence for encoding. Shaded areas denote standard error of the mean.

To perform a targeted investigation of how external and internal attention demands modulate memory brain state engagement, we extracted memory state evidence across the three phases (display: 0-500 ms, delay: 500-2500 ms, response: 2500-3500 ms) and performed three specific sets of analyses. Due to the structure of our mnemonic state classifier, positive values indicate more evidence for the memory retrieval state and negative values indicate more evidence for the memory encoding state.

We first sought to link the encoding state with external attention during the display phase. Specifically, we hypothesized that the encoding state would be recruited during initial stimulus presentation regardless of memory demands, as external attention demands should be equivalent across both perception and memory tasks. Alternatively, to the extent that the encoding state tracks memory formation either exclusively or in addition to external attention, display-interval encoding evidence should be greater for memory compared to perception trials. Following our pre-registration, we conducted a paired *t* -test comparing stimulus-locked memory state evidence across perception-display and memory-display trials (Figure 4A). We found significantly greater memory state evidence for memory-display (*M* = -0.0249, *SD* = 0.114) compared to perception-display trials (*M* = -0.0542, *SD* = 0.0916; *t* _38_ = 2.224, *p* = 0.0321, *d* = 0.2838). We separately compared memory state engagement within each task against zero via one-sample *t* -tests and found a significant decrease for perception-display trials, (*t* _38_ = -3.647, *p* = 0.0008, *d* = 0.5916) and not for memory-display trials (*t* _38_ = -1.344, *p* = 0.187, *d* = 0.218). Thus, we find significant engagement of the encoding state exclusively for the perception task. This result provides evidence against the alternative hypothesis that the encoding state tracks memory formation.

**Figure 4.**
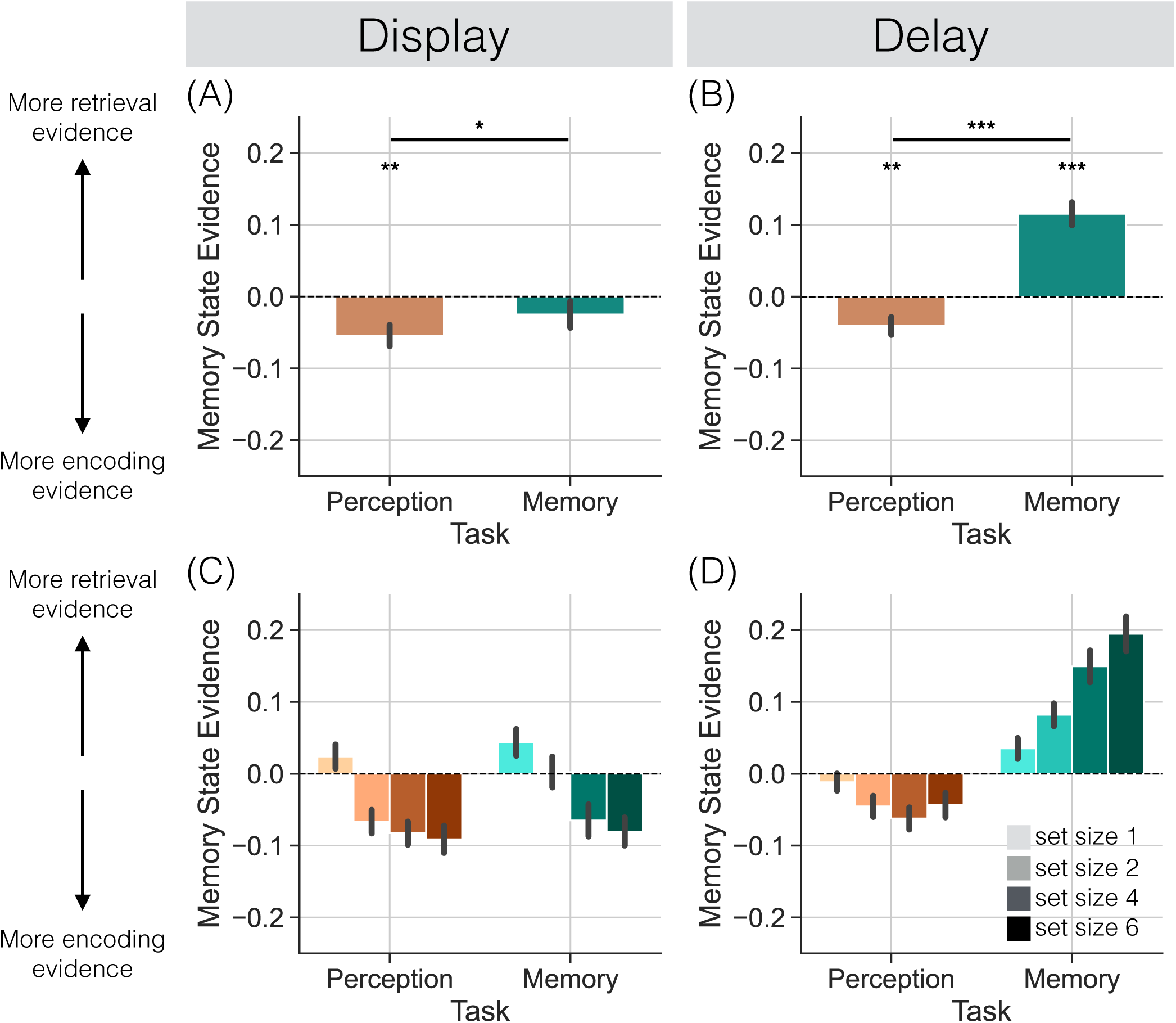
Memory state evidence by task phase. We assessed stimulus-locked memory state evidence as a function of task and set size. (A) Memory state evidence as a function of task during the display phase. There is a significant difference between memory state engagement during perception trials and memory trials (*p* = 0.0321). (B) Memory state evidence as a function of task during the delay phase. There is a significant difference between memory state engagement during perception trials and memory trials (*p <* 0.0001). (C) Memory state evidence as a function of task and set size during the display phase. There is a significant decrease in memory state engagement during perception and memory trials as a function of set size (*p <* 0.0001). (D) Memory state evidence as a function of task and set size during the delay phase. There is a significant increase in memory state engagement during memory trials as a function of set size, but no significant increase for perception trials (*p <* 0.0001). Error bars denote standard error of the mean. *p *<* 0.05; **p *<* 0.01; ***p *<* 0.001.

To further test the alternative hypothesis that the encoding state tracks memory formation, we assessed the extent to which memory state evidence during the display interval predicts subsequent task performance. Insofar as the encoding state tracks successful memory formation, greater display-phase encoding state evidence should selectively predict subsequent change detection accuracy. Display-phase encoding state evidence should not predict subsequent target detection performance given that attending to the squares will not facilitate target detection as the squares change location during the response phase. Following our pre-registration, we performed two regressions to relate memory state evidence to behavior. First, we conducted a linear regression with target detection RT as the dependent variable. We found no association between display-phase memory state evidence and RT (*β* = 3.824, *SD* = 17.30, *t* _38_ = 1.363, *p* = 0.1809, *d* = 0.2211). Second, we conducted a logistic regression with change detection accuracy as the dependent variable. We did not find a significant effect of display-phase memory state evidence on accuracy (*β* = 0.0389, *SD* = 0.4082, *t* _38_ = 0.588, *p* = 0.56, *d* = 0.0954). Having anticipated that memory state evidence would predict performance on the memory task, but not the perception task, we pre-registered a direct paired *t* -test to compare regression outputs between the perception and memory tasks. However, in view of our finding that neither task individually predicted behavior, we are unlikely to observe a difference between the two tasks, and we indeed did not find a significant difference between perception and memory (*β* = 0.0389, *SD* = 0.4082, *t* _38_ = 1.344, *p* = 0.1868, *d* = 0.3094). Together, these results show that memory state evidence does not predict subsequent performance in either task.

Our *a priori* expectation was that memory and perception trials would differ primarily during the delay phase. The logic is that the change detection (memory) task requires maintaining information over the delay, whereas the target detection (perception) task does not. Thus, to the extent that the retrieval state reflects internal attention, we should find more memory state evidence (equivalent to retrieval state evidence) for the delay phase of the memory compared to the perception task. This contrast provides the most robust assessment of maintenance-related internal attention as in both tasks no visual or external stimuli are present (excluding the fixation cross), thus the only difference is whether participants are (memory task) or are not (perception task) maintaining information during the delay.

Following our pre-registration, we conducted a paired *t* -test comparing stimulus-locked memory state evidence across perception-delay and memory-delay trials (Figure 4B). We find that memory state evidence is significantly greater for memory-delay (*M* = 0.1153, *SD* = 0.099) compared to perception-delay trials (*M* = -0.0408, *SD* = 0.0761; *t* _38_ = 7.305, *p <* 0.0001, *d* = 1.768). We separately compared memory state engagement within each task against zero via one-sample *t* -tests. For perception-delay trials, we find a significant decrease in memory state evidence (*t* _38_ = -3.301, *p* = 0.0021, *d* = 0.5355), indicating that participants engage the encoding state during the delay interval of the target detection task. For memory-delay trials, we find a significant increase in memory state evidence (*t* _38_ = 7.183, *p <* 0.0001, *d* = 1.165). These results show that the retrieval state is selectively engaged during the delay interval of the change detection task, but not the target detection task.

Having shown that working memory demands recruit the retrieval state during the delay interval, we next sought to link delay-phase memory state evidence to subsequent change detection performance. To the extent that the retrieval state tracks successful accessing of an internal representation, there should be a positive association between retrieval state evidence and subsequent memory performance. Alternatively, to the extent that retrieval state evidence tracks general internal attention and not memory success per se, there may be either no association or even a negative association between memory performance and retrieval state evidence. That is, greater retrieval state evidence may indicate a greater attempt to focus attention internally, reflecting difficulty rather than success. Following our pre-registration, we conducted a logistic regression with trial-level memory state evidence as the predictor and change detection accuracy as the outcome variable. Because we found an effect of set size on both behavior (above) and memory state evidence (below), we conducted four separate regression analyses, one for each set size, and averaged the resulting beta values together. The average beta value (*β* = 0.0807, *SD* = 0.8913) did not differ significantly from zero (*t* _38_ = 0.5578, *p* = 0.5802, *d* = 0.0905). These results show that greater retrieval state engagement does not predict successful subsequent performance in the change detection task.

To gain insights into the extent to which the retrieval state is modulated by decision making, we assessed response-locked memory state evidence during the response interval of each task. There are two alternative outcomes. First, to the extent that the retrieval state is recruited for any decision, we should find a gradual increase in memory state evidence (indicating greater retrieval state engagement) leading up to the response on both memory and perception trials. Alternatively, to the extent that the retrieval state is specifically recruited for a decision based on internal – rather than external – information, we should find a gradual increase in memory state evidence leading up to the response selectively for memory trials. Following our pre-registration, we conducted a 2 *×* 10 rmANOVA with factors of task and time interval (100 ms intervals from 500 ms pre-response to 500 ms post-response; Figure 5A). Response-locked memory state evidence during the response phase served as the dependent variable. We found a significant main effect of time (*F* _9,342_ = 64.05, *p <* 0.0001, 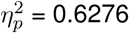) and task (*F* _1,38_ = 55.32, *p <* 0.0001, 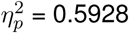), as well as a significant interaction between task and time (*F* _9,342_ = 5.359, *p <* 0.0001, 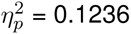).

**Figure 5.**
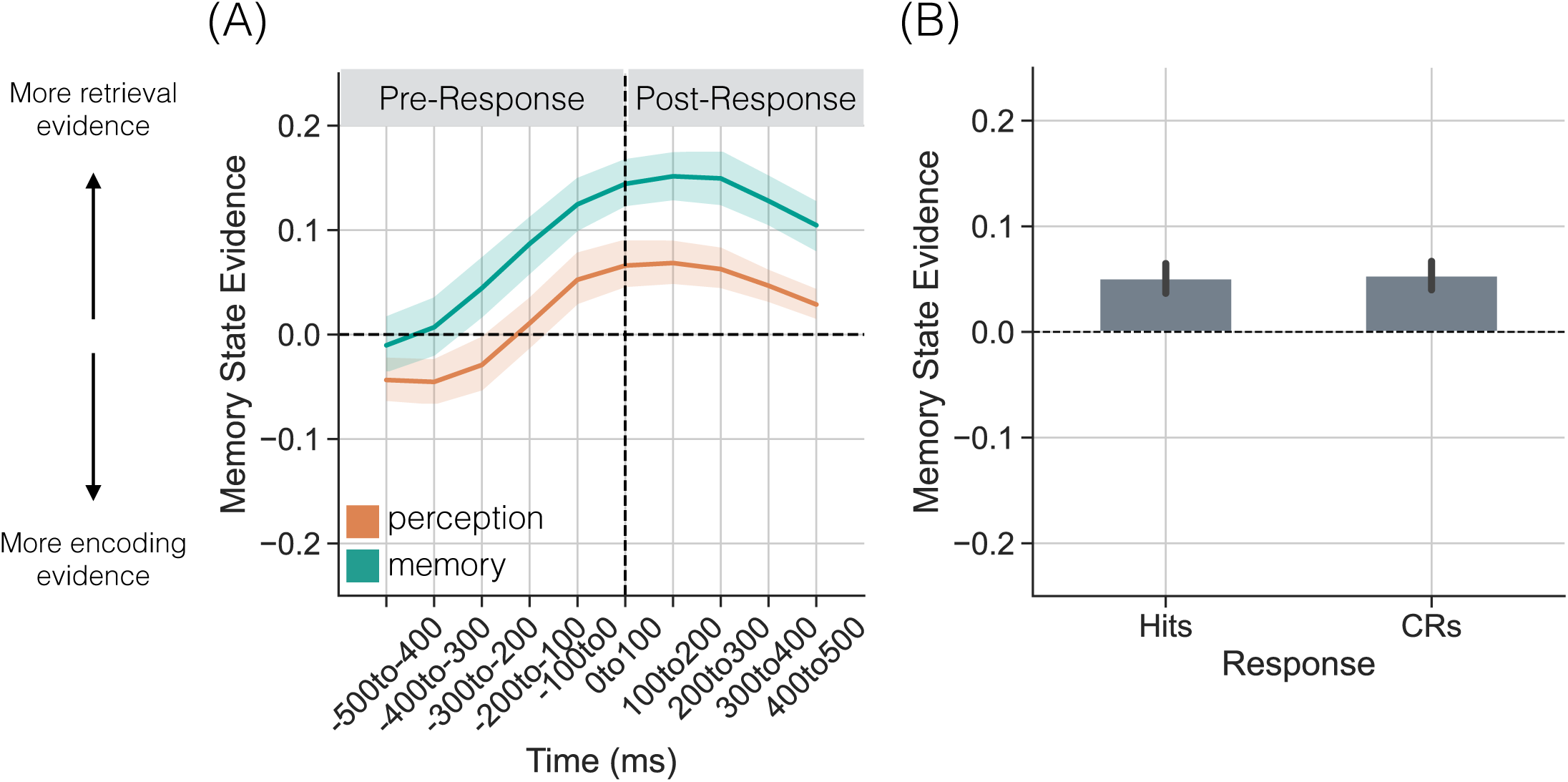
Response-locked memory state evidence. (A) Response-locked memory state evidence during the response phase. The bold dashed vertical line at time 0 to 100 ms indicates the onset of the response. (B) Response-locked memory state evidence as a function of memory response (hits or CRs) during the pre-response time interval (-500 to 0 ms). There is no significant difference between memory state engagement during hits and CR responses. Error bars denote standard error of the mean.

We anticipated a pre-response dissociation between memory and perception responses, thus, following our pre-registration, we conducted a post-hoc paired *t* -test to specifically compare memory state evidence between tasks during the pre-response (-500 to 0 ms) time interval. We found significantly greater memory state evidence for memory (*M* = 0.0505, *SD* = 0.0805) compared to perception responses (*M* = -0.0110, *SD* = 0.0660, *t* _38_ = 6.139, *p <* 0.0001, *d* = 0.8353). These results show that the retrieval state is specifically recruited for a decision based on internal – rather than external – information.

To further examine memory state engagement during the memory task and specifically during successful retrieval, we next assessed memory state evidence as a function of memory response (hit or correct rejection, CR) during the pre-response interval. To the extent that the retrieval state is engaged in the service of the attempt to retrieve a stored representation, we should find greater memory state evidence for CRs compared to hits, consistent with our prior work (Smith & Long, 2024; Wheelock & Long, 2024). Following our pre-registration, we averaged memory state evidence during the pre-response (-500 to 0 ms) time interval and used a paired *t* -test to compare memory state evidence across hit and CR trials in the memory task (Figure 5B). We did not find a significant difference between hit (*M* = 0.0507, *SD* = 0.0867) and CR trials (*M* = 0.0533, *SD* = 0.083; *t* _38_ = -0.3744, *p* = 0.7102, *d* = 0.0313). These findings suggest that correctly detecting that a square in the display has or has not changed colors recruits the retrieval state to a similar degree.

### Memory brain states are modulated by working memory load

We have shown that the demand to maintain information over a delay recruits the retrieval state. However, these findings do not address whether memory brain states track the direction of attention – external vs. internal – or the degree of internal attention. Put another way, the retrieval state may indicate whether attention is oriented internally rather than externally or the retrieval state may directly scale with the amount of internal attention that is deployed. By assessing memory state evidence as a function of set size, we can directly test these two possibilities.

We first tested the association between set size and display-phase memory state evidence. To the extent that increasing the amount of visual input recruits more external attention, we anticipated a positive association between set size and encoding state evidence. Following our pre-registration, we conducted linear regressions with factors of set size (1, 2, 4, 6) separately for perception and memory trials (Figure 4C). Stimulus-locked display-phase memory state evidence served as the dependent variable. We averaged the resulting beta values across both tasks and expected to find a significant negative beta, whereby memory state evidence decreases (reflecting greater encoding state engagement) as set size increases. We found a significant negative beta (*β* = -0.0222, *SD* = 0.0145, *t* _38_ = -9.399, *p <* 0.0001, *d* =1.525). Given our hypothesis that the encoding state tracks external attention as opposed to successful memory formation, we expected that the association between display-phase memory state evidence and set size would not differ between perception and memory trials. We conducted a paired *t* -test and did not find a significant difference in beta values for perception vs. memory task trials (*t* _38_ = 1.571, *p* = 0.1244, *d* = 0.3191). Thus, as more visual information is presented, encoding state evidence increases, suggesting that this state reflects external attention to environmental stimuli.

As we demonstrated above, working memory demands during the delay phase strongly recruit the retrieval state. The critical question is whether recruitment of the retrieval state is further modulated by working memory load or the amount of information that participants must maintain. To the extent that the retrieval state tracks the degree of internal attention, we expect to find greater memory state evidence as working memory load increases. Following our pre-registration, we conducted a linear regression with factors of set size (1, 2, 4, 6) selectively for the memory trials (Figure 4D). Stimulus-locked delay-phase memory state evidence served as the dependent variable. We expected to find a significant positive beta, whereby memory state evidence increases (reflecting greater retrieval state engagement) as set size increases. We found a significant positive beta (*β* = 0.0316, *SD* = 0.0277, *t* _38_ = 7.021, *p <* 0.0001, *d* = 1.139). To demonstrate that this positive association between set size and memory state evidence is specific to the memory task, we directly compared the regression outputs across perception-delay and memory-delay trials. We found a significantly greater beta for the memory task trials compared to perception task trials (perception: *β* = -0.0057, *SD* = 0.0180; *t* _38_ = 6.662, *p <* 0.0001, *d* = 1.593). These results show that memory state evidence increases (reflecting greater retrieval state engagement) as set size increases selectively for the memory task.

## Discussion

The aim of this study was to directly connect memory brain states with the external/internal axis of attention. Using cross-study decoding of scalp EEG data, we measured memory state engagement during a working memory (WM) paradigm in which participants performed either target or change detection. We find that the encoding state is recruited during the display interval of both target and change detection tasks, whereas the retrieval state is selectively engaged during the delay interval of the change detection task – and modulated by set size – but not the target detection task. Together, these results suggest that memory states map onto the external/internal axis of attention to support both working and long-term memory.

We find that memory brain states are modulated by the different phases – display, delay, response – of a WM paradigm. According to prior work, the retrieval state specifically enables intentional retrieval from long-term episodic memory (Tulving, 1983; Rugg & Wilding, 2000), in which case it should not be modulated in the current paradigm as there were no demands to access long-term memories associated with a particular spatiotemporal context. However, we find engagement of both memory encoding and memory retrieval states across the display, delay, and response intervals and across tasks. These results indicate that memory brain states track processes related to working memory, including external and internal attention.

During the display phase of both the target and change detection tasks, we find encoding state engagement. To the extent that the encoding state tracks memory formation, it should be strongly and selectively engaged during the display phase of the change detection task, as encoding the display is needed to later report on color changes. However, we find similar levels of encoding state engagement across both tasks and that encoding state evidence increases as a function of set size. Attention is recruited when stimuli compete (Desimone & Duncan, 1995) and more stimuli will lead to more competition (Kastner, Pinsk, De Weerd, Desimone, & Ungerleider, 1999). Thus, encoding state engagement may reflect an attentional response to increasing perceptual demands as the number of stimuli increases. Together, these findings support the interpretation that the encoding state tracks external attention to stimuli in the environment. We anticipate that the encoding state would be recruited in support of visual search (Treisman & Gelade, 1980; Wolfe, 1998) and may play a role in change blindness and the attentional blink (Rensink, O’Regan, & Clark, 1997; Shapiro, Raymond, & Arnell, 1997).

We find robust retrieval state engagement selectively during the change detection task delay interval. This finding provides strong support for the hypothesis that the retrieval state reflects internal attention. WM maintenance can be considered a form of internal attention (Chun, 2011; Kiyonaga & Egner, 2016; Vo et al., 2021, c.f. Myers, Stokes, & Nobre, 2017), whereby attention is directed internally to no longer perceptually present information. Internal attention relies on neural substrates common to memory retrieval, including the default mode network (DMN; Smallwood et al., 2013; H. Kim, 2015; Buckner & DiNicola, 2019) and spectral signals in the theta and alpha bands (Kam et al., 2019; Woodman et al., 2022). Our interpretation is that a single, internal attention brain state – here labeled the retrieval state – tracks all of these processes, from long-term episodic memory retrieval, to WM maintenance, to other forms of internal mentation such as mind wandering (Long, 2023; Buckner & DiNicola, 2019). Interestingly, there is some retrieval state engagement during the target detection task delay interval, at least within the first several hundred milliseconds. It may be that some degree of internal attention is invoked when external input is removed, regardless of intentional maintenance demands. Such a finding is reminiscent of prior work showing increased DMN activity during ‘idling’ or mind-wandering in the absence of visual input (Mason et al., 2007) and increases in alpha power for eyes-closed compared to eyes-opened conditions (Geller et al., 2014). These transitions between brain states may be indicative of input-driven switches between external and internal states (Verschooren, Pourtois, & Egner, 2020; Nobre & Gresch, 2025) and are an important avenue for future work.

The retrieval state is selectively modulated by WM load. As the amount of information to-be-maintained increases, retrieval state engagement increases, but only during the change detection task. Thus, it is not merely that internal attention is recruited in the absence of visual input, but directly scales with the amount of maintained information. Previous work has shown that contralateral delay activity (CDA), a negative voltage deflection over the hemisphere contralateral to the visual hemifield in which information is maintained, increases up to WM capacity limits (Vogel & Machizawa, 2004; Feldmann-Wüstefeld, Vogel, & Awh, 2018). Thus the CDA is thought to track specific objects maintained in WM (Hakim, Adam, Gunseli, Awh, & Vogel, 2019). As we observe maximal retrieval state engagement with the largest set size (6), which is beyond the average capacity limit (Luck & Vogel, 1997; Cowan, 2010), it is likely that the retrieval state tracks internal attention engaged in service of the *attempt* to maintain information and may not directly relate to the items themselves. To the extent that the retrieval state and WM maintenance invoke the same underlying alpha power mechanism (Jensen et al., 2002; Cooper et al., 2003; Vo et al., 2021), the retrieval state could reflect a map of “prioritized space” (Hakim, Awh, & Vogel, 2021). This interpretation is consistent with our prior work in which evidence for the retrieval state is positively associated with maintained information in a variant of the Posner spatial cueing task (Posner, 1980; Y.-H. Kim et al., 1999; Long, 2023). Thus, the retrieval state likely reflects a domain-general internal attention signal.

Finally, we show that the retrieval state is recruited while participants detect changes and targets, with greater recruitment for change detection. As with prior long-term episodic memory retrieval work (Smith & Long, 2024), we speculate that retrieval state engagement during the response phase reflects internal attention directed to stored representations, in this case, the immediately preceding display interval stimulus array. We have previously observed greater retrieval state evidence for correct rejections compared to hits (Smith & Long, 2024; Wheelock & Long, 2024) which we interpreted as reflecting an internal search and/or evidence accumulation process (e.g. Ratcliff & McKoon, 2008) used to reject not-studied lure stimuli. That we failed to find a dissociation between correct rejections and hits in the current study is instead consistent with the view that participants perform an internal comparison process (Sternberg, 1966) between the test probe and the contents of their WM to detect both changed and unchanged displays.

Taken together, our findings provide strong evidence that memory encoding and memory retrieval states map onto the external/internal axis of attention. The degree to which memory states reflect the orientation of attention has important consequences for our understanding of memory and attention as well as cognition more broadly. Although our results demonstrate an overlap between memory brain states and attention, a direct investigation of the shared and unique components is necessary in order to constrain our conceptualization of the two domains of ‘attention’ and ‘memory’ (Logan et al., 2021). Our results suggest that attentional processes may contribute substantially to memory success and are consistent with recent work showing that a composite of attention information predicts subsequent memory (Mirjalili & Duarte, 2025). The current line of work on memory brain states stems from theories and data showing that memory states are governed by hippocampal circuitry (Hasselmo, Bodelón, & Wyble, 2002; Hasselmo, 2005). Thus, a critical open question is how the brain states currently indexed relate to hippocampal signals. To the extent that the hippocampus sits atop the sensory/perceptual hierarchy (Aly & Turk-Browne, 2016; Turk-Browne, 2019; Nobre & Gresch, 2025), hippocampal signals may impact the currently observed cortical brain states. Alternatively, there may be memory-specific signals, distinct from the current states, driven by the hippocampus. For instance, lingering hippocampally-mediated memory states (Duncan, Ketz, Inati, & Davachi, 2012) may be induced by judgments made about old and new stimuli whereas cortical states may be invoked to re-orient attention when a task-irrelevant hippocampal state is engaged (Wheelock & Long, 2024). More broadly, brain states govern downstream processing (Long & Kuhl, 2021) and impact behavior (Harris & Thiele, 2011; Kay & Frank, 2019). To the extent that the current cortical states track attention vs. memory processes will result in different influences on behavior. Whereas retrieval trades off with encoding (Hasselmo et al., 2002; Long & Kuhl, 2019), meaning that retrieval will prevent encoding the current moment and impair subsequent memory (Long & Kuhl, 2019), internal focus may facilitate memory by enabling elaborative, constructive internal processes such as imagery (Craik & Lockhart, 1972; Craik & Tulving, 1975). By identifying a core set of attentional states, our work will enable online, real-time detection of brain state engagement, identification of which states are optimal based on task goals and desired behaviors, and the development of tools to correct maladaptive or suboptimal state engagement.

In conclusion, we demonstrate that memory encoding and memory retrieval states map onto the external/internal axis of attention, respectively. We find that the retrieval state is selectively engaged when there is a demand to maintain and is further modulated by memory load. Our results show that memory brain states are recruited in response to attentional demands in support of both working and long-term memory. Together, these results advance our understanding of how attention and memory processes overlap, informing our conceptual framing and empirical measurement of how these processes impact cognition.

**Table 1.**
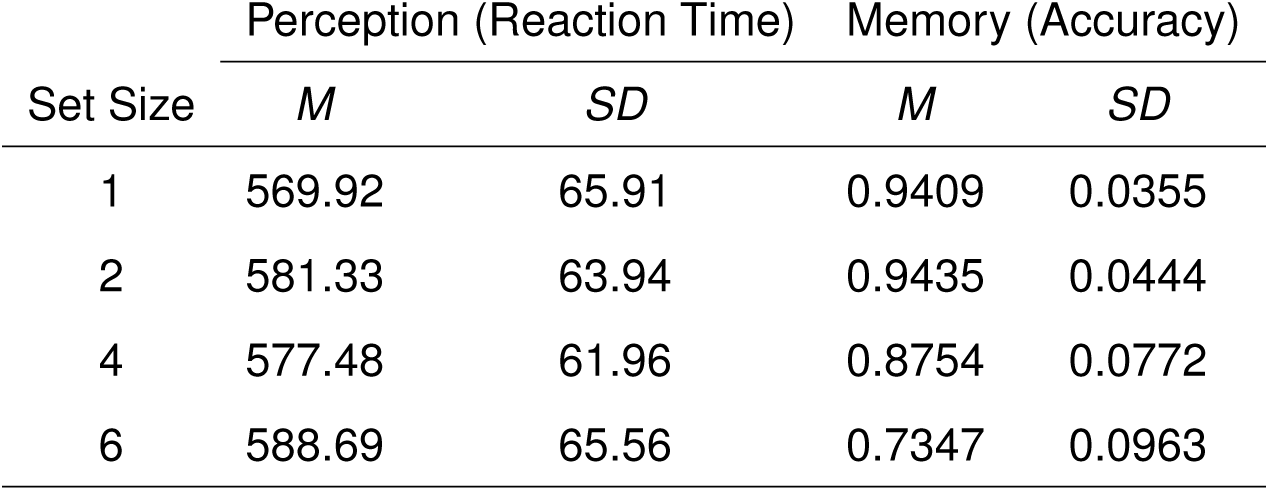
Behavioral performance as a function of task and set size.

| Set Size | Perception (Reaction Time) |  | Memory (Accuracy) |  |
| --- | --- | --- | --- | --- |
|  | *M* | *SD* | *M* | *SD* |
| 1 | 569.92 | 65.91 | 0.9409 | 0.0355 |
| 2 | 581.33 | 63.94 | 0.9435 | 0.0444 |
| 4 | 577.48 | 61.96 | 0.8754 | 0.0772 |
| 6 | 588.69 | 65.56 | 0.7347 | 0.0963 |

## Acknowledgments

This work was supported by a grant from the National Institutes of Health (NINDS R01 NS132872, PI: N.M.L.).

